# Expression of a tetracycline-controlled transactivator (Tet-On/Off system) in beta cells reduces insulin expression and secretion in mice

**DOI:** 10.1101/2021.02.10.430692

**Authors:** Nathalie Jouvet, Khalil Bouyakdan, Cindy Baldwin, Jadwiga Marcinkiewicz, Thierry Alquier, Jennifer L. Estall

**Affiliations:** Institut de recherches cliniques de Montreal (IRCM), Montreal, Quebec, Canada; University of Montreal, Montreal, Quebec, Canada; Centre de Recherche du CHUM, Université de Montréal, Quebec, Canada; Montreal Diabetes Research Centre, Université de Montréal, Quebec, Canada

**Keywords:** β cell, tetracycline-controlled transactivator, tTA, MIP, TetO-Cre, insulin expression

## Abstract

Controllable genetic manipulation is an indispensable tool in research, greatly advancing our understanding of cell biology and physiology. However, in beta cells, transgene silencing, low inducibility, ectopic expression and off-targets effects on cell function and glucose homeostasis are a persistent challenge. In this study, we investigated whether an inducible, Tet-Off system with beta-cell specific MIP-itTA driven expression of TetO-Cre^Jaw/J^ could circumvent previous issues of specificity, efficacy and toxicity. Following assessment of tissue-specific gene recombination; beta cell architecture; *in vitro* and *in vivo* glucose-stimulated insulin secretion (GSIS); and whole-body glucose homeostasis, we discovered that expression of any tetracycline-controlled transactivator (e.g. itTA, rtTA or tTA) in beta cells significantly reduced *Insulin* gene expression and decreased insulin content. This translated into lower pancreatic insulin levels and reduced insulin secretion in mice carrying a MIP-itTA transgene, independent of Cre-recombinase expression or doxycycline treatment. These results raise significant concern regarding the use of Tet-On or Tet-Off systems for genome editing in beta cells and emphasize the need to control for effects of transactivator expression. Our study echoes ongoing challenges faced by fundamental researchers focused on beta cells and highlights the need for consistent and careful control of experiments using these research tools.

## INTRODUCTION

Mice are an invaluable research model leading to numerous translatable discoveries related to human drugs and greatly expanding our understanding of normal physiology and disease. Transgenic mouse models relying on cell-specific promoters, with or without inducible control, allow reliable over-expression or ablation of genes of interest in a temporal and spatial matter. Popular strategies to control transgene expression include the estrogen receptor tamoxifen-dependant ligand domain (ERT)^1^ and the Tet Operon(TetO)/repressor bi-transgenic systems^2-4^. Cell-specific and drug-controlled transgene expression systems are now widely used and there are countless mouse models with tissue-specific promoters available for mice.

There have been numerous challenges with targeting transgene expression to pancreatic beta cells^5^. Multiple promoters can achieve cell-type specificity; however ectopic expression, primarily in the brain, is suspected to be responsible for reported off-target effects involving reduced beta cell function and altered glucose homeostasis^6,7^. The once popular rat insulin promoter (*RIP* or *Ins2*) has undeniable efficiency^8^, but can drive expression in several regions of the central nervous system (CNS) and impact glucose tolerance^6,7,9^. The *Pdx-1* (pancreatic and duodenal homeobox 1) promoter, also known as insulin promoter factor 1 (*Ipf1*), is a useful tool, as it can drive expression in the whole pancreas or only beta cells depending on the timing of promoter activation^10^. However, the *Pdx-1* promoter also promotes expression in other areas of the digestive track^10^ and the CNS under some circumstances^6,11^. In contrast, the mouse insulin promoter (*MIP* or *Ins1*) has not yet been shown to drive ectopic transgene expression in the brain or other neuroendocrine cell types^6,12,13^; however, efficiency of one widely available model using an inducible variant (Ins1-CreERT^Thor^) appears highly variable^14,15^. There are new concerns surrounding off-targets effects of certain transgene elements, such as the mini-gene for human growth hormone (hGH) often included in transgenic constructs to enhance transcriptional efficiency^16,17^. Unanticipated hGH protein expression from this element is now implicated in a number of problems linked to various transgenic models and ^18-21^ can have significant effects on beta cell function and islet mass^12,17^.

In addition to cell specificity and transgene design issues, exogenous drugs used to control transgene expression can cause toxicity. Tamoxifen can impair glucose tolerance in a strain and sex specific manner^22^, alter beta cell function^17^ or proliferation^23^, and requires long wash out periods^24^. In recent years, mutations added to the original ERT sequence increased its effectiveness and lowered the required dose of tamoxifen^25^ but doses necessary to achieve desired recombination efficiency can still have lasting negative effects^15^. The tTA/Tet Operon system to control transgene expression is an alternative to ERT/tamoxifen. Models using *Ins1*-or *Ins2*-driven reverse tetracycline-driven transactivator (rtTA) expression have been successfully used to control transgene expression in beta cells^26,27^; however, like the ERT system, the Tet-On system relies on drug treatment during the time of experimentation. Tetracycline/Doxycycline can be toxic to mitochondrial function^28-30^; and can independently impact glucose homeostasis in diabetic *db/db* mice^31^.

A benefit of the tTA/TetO system is its versatility to drive expression of *any* transgene downstream of TetO. Although resistance to using this system stems from the need to have two to three transgenic mouse lines to generate test groups, the Tet-Off system avoids use of exogenous drugs when experimental outcomes are being measured. The improved tTA transactivator (itTA)^32^ has been successfully used in combination with the MIP promoter to drive islet transgene expression^33^. However, in light of recent challenges in other models^14,15,31,34,35^, additional studies are required to evaluate the efficiency, specificity, and potential off-target effects of this system for beta cell targeting.

Using a combination of the Tet-Off system and the MIP (*Ins1*) promoter (MIP-itTA^33^:TetO-Cre^Jaw/J36^), we characterized this new mouse line in terms of recombination efficiency and cell-type specificity. We also evaluated effects of each transgene (separate and together) on insulin secretion in response to glucose. Our data show that tetracycline transactivators can have significant negative effects on insulin expression and beta cell function, and that leaky expression of the TetO operon reduces cell-type specificity in this system. These data highlight important limitations of the Tet-On/Off model systems for cell-specific targeting, particularly in beta cells.

## RESEARCH DESIGN AND METHODS

### Generation of mice

Unless specified, hemizygote male and female MIP-itTA^33^ or TetO-Cre^Jaw/J36^ (B6.Cg-Tg(tetO-cre)1Jaw/J, stock no 006234) mice on a mixed C57Bl/6J:C57Bl/6N background were used for all experiments. To generate reporter mice, transgenic MIP-itTA or TetO-Cre^Jaw/J^ mice were crossed with homozygous mT/mG^37^ mice on a mixed C57Bl/6J:129Sv:CD-1 background. Control mice were age- and sex-matched littermates. All mice (unless indicated) received doxycycline (Bioshop) in the drinking water (0.1 g/L to mothers during breeding, pregnancy and lactation or 1 g/L to pups after weaning) until they were 8 weeks of age. A minimum 5-week wash-out period following doxycycline was provided. Mice were maintained on a 12-hour dark/light cycle and given free access to water and standard laboratory chow (Teklad diets 2018). Mice were maintained and sacrificed according to approved protocols from the Institut de recherches cliniques de Montreal (IRCM). Genotyping of tail DNA was performed using primers listed in Supplementary Table 1.

### Gene expression analysis

RNA extracted by Trizol (Invitrogen) or RNeasy Mini Kit (Qiagen) from INS-1 cells and islets (∼120), respectively, was reverse transcribed (Life technologies) and mRNA measured by qPCR using SYBR green (Bioline). Relative expression was calculated by ΔΔct-method and normalized to TATA-box binding protein (*Tbp*). Primers are listed in Supplementary Table 1.

### Histological analysis of tissues in mT/mG reporter mice and immunohistochemistry

At 18-20 weeks of age, male mice without doxycycline were perfused intracardially with 4% paraformaldehyde under anaesthesia. The brains were post-fixed 3 h in 4% paraformaldehyde, cryopreserved in 20% sucrose, and embedded in OCT (VWR) and cryosectionned at 18 μm using a cryostat (CM3050s Leica, Wetzlar, Germany). Sections were mounted and imaged with a Zeiss fluorescent microscope (Axio Imager M2 Carl Zeiss AG, Jena, Germany). All other tissues were frozen in OCT using isopentane (Fisher) and kept at -80°C until sectioning. For whole body sectioning, 10 day old pups were euthanized by CO_2_ and immersed in 2-Butanol (Fisher) and kept at -80°C until sectioning. Tissues were sectioned (7-10 μm) using a Leica cryostat. Paraffin-embedded pancreas were cut in sections of 5 μm and stained with Haematoxylin and Eosin (H&E). Immunofluorescence of paraffin sections was performed using guinea pig anti-insulin (1:100, Dako), mouse anti-glucagon (1:250, Sigma) antibodies and the corresponding secondary antibodies Alexa-488 or Cy3 (Jackson Immunoresearch) with ProLong Gold Antifade Mountant with DAPI (ThermoFisher Scientific). The remaining immunofluorescence images were taken using a Leica DM5500B microscope at a magnification of 20X for islets and 10X for other tissues and analyzed with Volocity software. Whole mount images of mice were taken using a Leica DM6 and a mosaic of multiple small images were assembled using LASX software. Confocal images of the intestines were taken with a ZEISS LSM700 at a magnification of 20X and analyzed with ZEN software.

### Cell culture

INS-1 cells were cultured in RPMI 1640 medium (Wisent, 11.1 mM glucose) supplemented with 10% (vol/vol) heat-inactivated FBS (Wisent), 2-mercaptoethanol (50 μM, Bioshop), sodium pyruvate (1 mM, Wisent), L-glutamine (2 mM, Wisent), HEPES (10 mM, Bioshop) and penicillin/streptomycin (Wisent). 293FT cells were maintained in DMEM (Wisent, 25 mM glucose) supplemented with 10% FBS and penicillin/streptomycin. Cells were grown in 5% (vol/vol) CO_2_ at 37°C and passaged at ∼70% confluence. Culture medium was changed every 2–3 d.

### Production of Lentiviruses, Transduction of Stable Clones and Insulin Content

In the vector pLJM1-EGFP (Addgene #19319^38^), EGFP was replaced by itTA (from pBSitTA-nls, gift from Rolf Sprengel^32^), rtTA (Addgene #25434^39^) or tTA (Addgene #14901^40^) by Gibson cloning (NEB) or was ligated without GFP (empty vector). Transferring plasmids psPAX2 (Addgene #12260) and pMD2.G (Addgene #12259) were cotransfected with each pLJM1 constructs. For transduction, INS-1 were plated in 100 mm plates at the density of 3 × 10^6^ cells/plate and cultured overnight. Cells were transduced in basic media (without FBS or antibiotics) for 8 hours (≈2 × 10^7^ TU/ml for each lentivirus, MOI of 0,1) and stable clones selected 36 hours later using 0.5 μg/ml of puromycin (Biobasic). Each stable clone was plated at a density of 3 × 10^5^ cells/ml in a 24 well plate and kept overnight. The next day, cells were washed in PBS and 1 ml of acid–ethanol (1.5% HCl in 70% EtOH) was added and cells incubated at -20°C for 24 hours. Samples were neutralized with 1 M Tris pH 8.0 and insulin was measured using the STELLUX Chemiluminescent Rodent Insulin ELISA kit (Alpco) and content normalized to DNA concentration (Biobasic).

### Islet isolation

Mouse pancreas was digested following perfusion with 0.4 U/ml Liberase TL (MilliporeSigma) in Hank’s Balanced Salt Solution (HBSS, Wisent) buffer with Ca^2+^/Mg^2+^ and dispersed by gentle shaking in buffer without Ca^2+^/Mg^2+^ containing 0.1% BSA and 20 mM HEPES, pH 7.4. Pellet was washed in HBSS buffer and resuspended in Histopaque-1077 (Sigma) overlaid with RPMI 1640 (without glucose) prior to separation by centrifugation. Handpicked islets were cultured overnight in 11 mM glucose RPMI 1640 (10% FBS and penicillin/streptomycin).

### Static *in vitro* insulin secretion

Insulin secretion was assessed in 1-h static incubations of isolated islets as previously described^12^. Briefly, batches of ten islets each were washed twice for 20 minutes in KRBH solution containing 0.1% (wt/vol.) BSA and 2.8 mmol/l glucose. Islets from at least 2 different mice (in 12 replicate batches) were incubated at 37°C for 1 h in the presence of 2.8 mmol/l or 16.7 mmol/l glucose (with and without KCl). Intracellular insulin content was measured after acid–ethanol (1.5% HCl in 70% EtOH) incubation overnight at -20°C. Samples were neutralized with 1 M Tris pH 8.0 before insulin was measured with the mouse insulin ELISA immunoassay kit (Alpco).

### Glucose/insulin tolerance tests and serum insulin

Physiological tests were performed at 13-18 weeks of age. After a 16 h-fast, glucose tolerance tests (GTTs) were performed following oral gavage (OGTT) or intraperitoneal (IPGTT) injection of glucose diluted in water (1.5 g/kg). Areas under the curve (AUC) were calculated using baseline fasting glucose. Insulin tolerance tests were performed following intraperitoneal injection of human insulin (Humulin R, Lilly, 0.8/kg) in 4 h-fasted mice. Glucose was measured from the tail vein by glucometer (FreeStyle Lite, Abbott Diabetes Care) or in serum using the mouse ultrasensitive insulin ELISA (Alpco).

### Total pancreatic insulin content

25-35% of the pancreas (region closest to the spleen) was incubated overnight at -20°C in 5 ml acid-ethanol (1.5% HCl in 70% EtOH). Tissue was then homogenized in the acid-ethanol and homogenate further incubated overnight at -20°C. Homogenate was centrifuged for 15 min at 2000 rpm (4°C) prior to neutralization of supernatant with 1 M Tris pH 8.0. Insulin was measured by STELLUX Chemiluminescent Rodent Insulin ELISA kit (Alpco) and content normalized to initial tissue weight.

### Statistical Analysis

Data comparing two or more groups on one variable were analyzed by one-way ANOVA, corrected for multiple comparisons (Dunnett). For data sets with two variables, two-way ANOVA was used, corrected for multiple comparisons (Dunnett). Analyses were performed using GraphPad Prism. Unless indicated, values are mean ± SEM.

## RESULTS

### Inducible Cre-recombinase is efficiently targeted to islets of MIP-itTA:TetO-Cre^Jaw/J^ mice

To design a Tet-Off-based mouse model expressing Cre-recombinase only in beta cells, we paired two existing models: the MIP-itTA^33^ and the TetO-Cre^Jaw/J 36^. In this system, tetracycline or its derivatives (e.g doxycycline) represses expression of TetO-driven Cre-recombinase by inhibiting binding of the tetracycline transactivator (itTA) to the Tetracycline Operator sequence (TetO). When doxycycline is removed, the transgene is expressed. Given that a number of transgenic mouse models have off-target effects due to the human growth hormone (hGH) minigene^21^, we confirmed that this element was not present in MIP-itTA or TetO-Cre^Jaw/J^ genomes by PCR using hGH minigene specific primers (Figure 1A).

**Figure 1:**
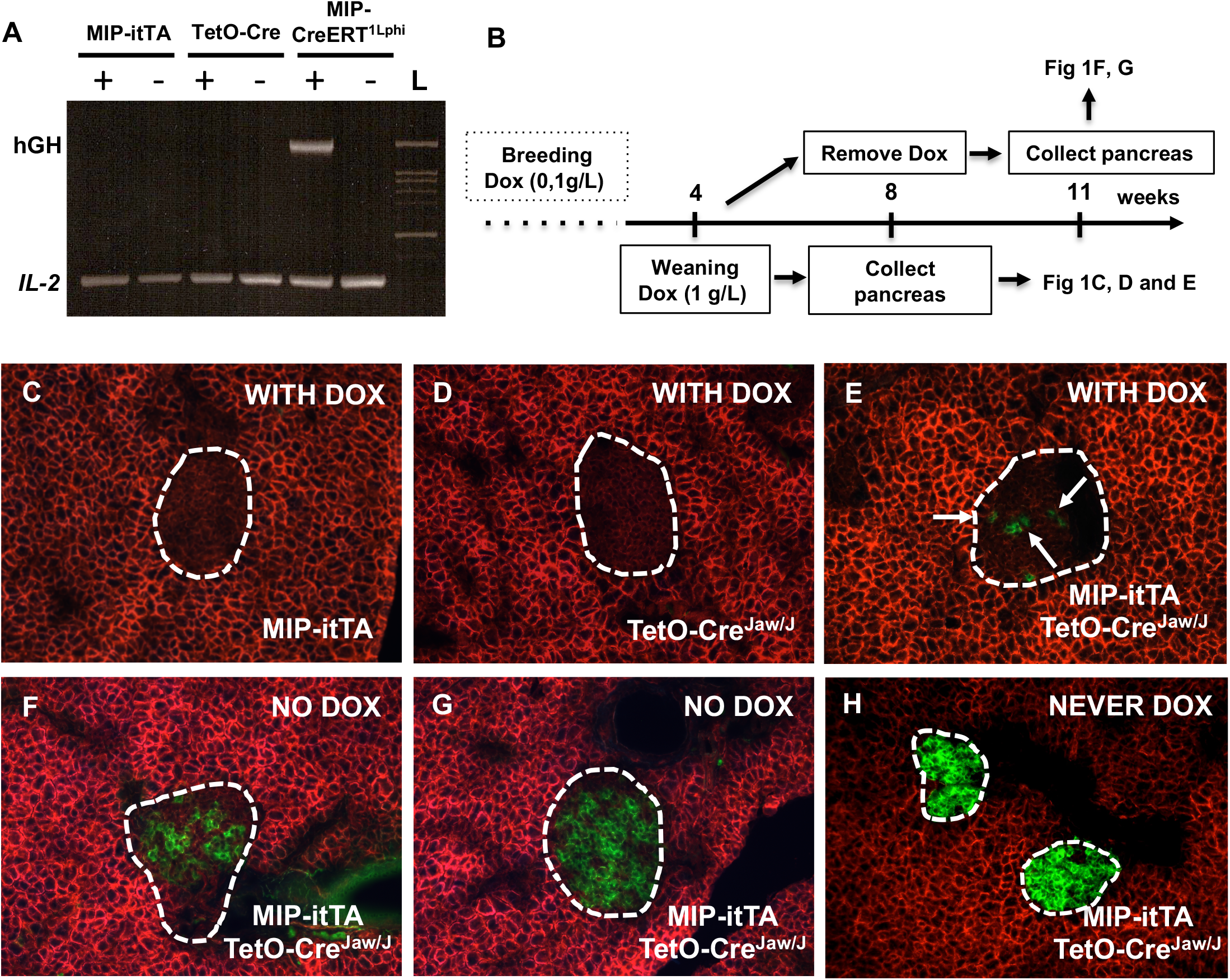
Cre recombination validated in islets of MIP-itTA:TetO-Cre^Jaw/J^ mice. *A*: PCR analysis to detect the human growth hormone (hGH) minigene in genomic DNA from MIP-itTA or TetO-Cre^Jaw/J^ mice. Tail DNA from MIP-CreERT^Lphi^ and from WT littermate mice were used as positive and negative controls, respectively. *B*: Schematic of the experimental design and timeframe. Pancreatic sections from *C*: MIP-itTA:mTmG, *D*: TetO-Cre^Jaw/J^:mTmG or *E*: MIP-itTA:TetO-Cre^Jaw/J^:mTmG mice kept on doxycycline treatment (*in utero* and 8 weeks after birth, WITH DOX) or *F-G*: 13-week-old MIP-itTA:TetO-Cre^Jaw/J^:mTmG mice without doxycycline for the last 4 weeks (NO DOX). *H*: Pancreatic sections from MIP-itTA:TetO-Cre^Jaw/J^:mTmG mice never exposed to doxycycline (NEVER DOX) (representative images of *n* = 2 mice, male or female).

To test the efficiency of recombination in beta cells, we crossed MIP-itTA:TetO-Cre^Jaw/J^ mice with the mTmG reporter mouse line^37^. In this model, all cells express red fluorescent protein (RFP) in the absence of Cre-recombinase (transgene expression is off in the presence of doxycycline). After washout of doxycycline, the transactivator can bind to the Tet operon and drive expression of Cre-recombinase, excising the RFP gene and replacing it with green fluorescent protein (GFP). All mice were given doxycycline (dox) in the drinking water from initiation of mating until 8 weeks of age (Figure 1B). At 8 weeks of age, one group continued DOX treatment (WITH DOX groups, Figures 1C-E). In a separate group, doxycycline was withdrawn, and mice were kept for an extra 4 weeks to monitor Cre-mediated recombination (NO DOX group, Figure 1F-G). As expected, mice lacking one transgene of the complete Tet system (transactivator or Tet operon) showed no GFP-positive cells, demonstrating lack of inappropriate Cre-recombinase expression in islets (i.e. MIP-itTA:mTmG, Figure 1C or TetO-Cre^Jaw/J^:mTmG mice, Figure 1D). When the system was complete (MIP-itTA:TetO-Cre^Jaw/J^:mTmG), we observed a few green cells in mice receiving doxycycline (which should prevent transgene expression), representing a low level of “leaky” itTA activity (Figure 1E, white arrows). Of all the islets analysed for the study (about 45), we estimate that about 2-5% of the cells expressed Cre-recombinase in the presence of doxycycline. In contrast, mice taken off doxycycline for 4 weeks had high levels of GFP-positive beta cells in the islets with about 50-90% efficiency (Figure 1F-G are representative images of *F*:low and *G*:high recombination), indicating that removal of doxycycline led to appreciable increases of Cre-recombinase expression in islets. Finally, we included a group of mice that never had doxycycline in the drinking water (NEVER DOX, Figure 1H) where we observed close to 100% recombination efficiency in all the islets.

### TetO-Cre^Jaw/J^ mice exhibit ectopic Cre-recombination in regions of the brain and the digestive system

To determine cell-type specificity of transgene expression, we tested for the presence of Cre-recombinase in multiple other tissues. In the central nervous system of adult mice expressing only the TetO-Cre transgene (TetO-Cre^Jaw/J^:mTmG, Figure 2A-D and Supplemental Figure 1A-D for controls), we observed no GFP-positive cells in the arcuate nucleus, paraventricular nucleus or nuclei of the brain stem (Figure 2A-C), but we did find a subset of GFP positive cells in the choroid plexus (Figure 2D, white arrows). We did not detect GFP-positive cells in the liver, muscle or heart (Figure 2E-G, controls in Supplemental Figure 1E-G). However, we noted many green cells in the stomach (Figure 2H, white arrow) and the gastrointestinal wall of the duodenum (Figure 2I), white arrow) and a few green cells in other sections of the intestine compared to control littermates (Figure 2J-L, controls in Supplemental Figure 1H-L). Confocal analysis of the duodenum and ileum confirmed the presence of Cre-recombinase mediated recombination in the submucosa (Figure 2M-N) and in the villi of the jejunum and colon (Figure 2O-P).

**Figure 2:**
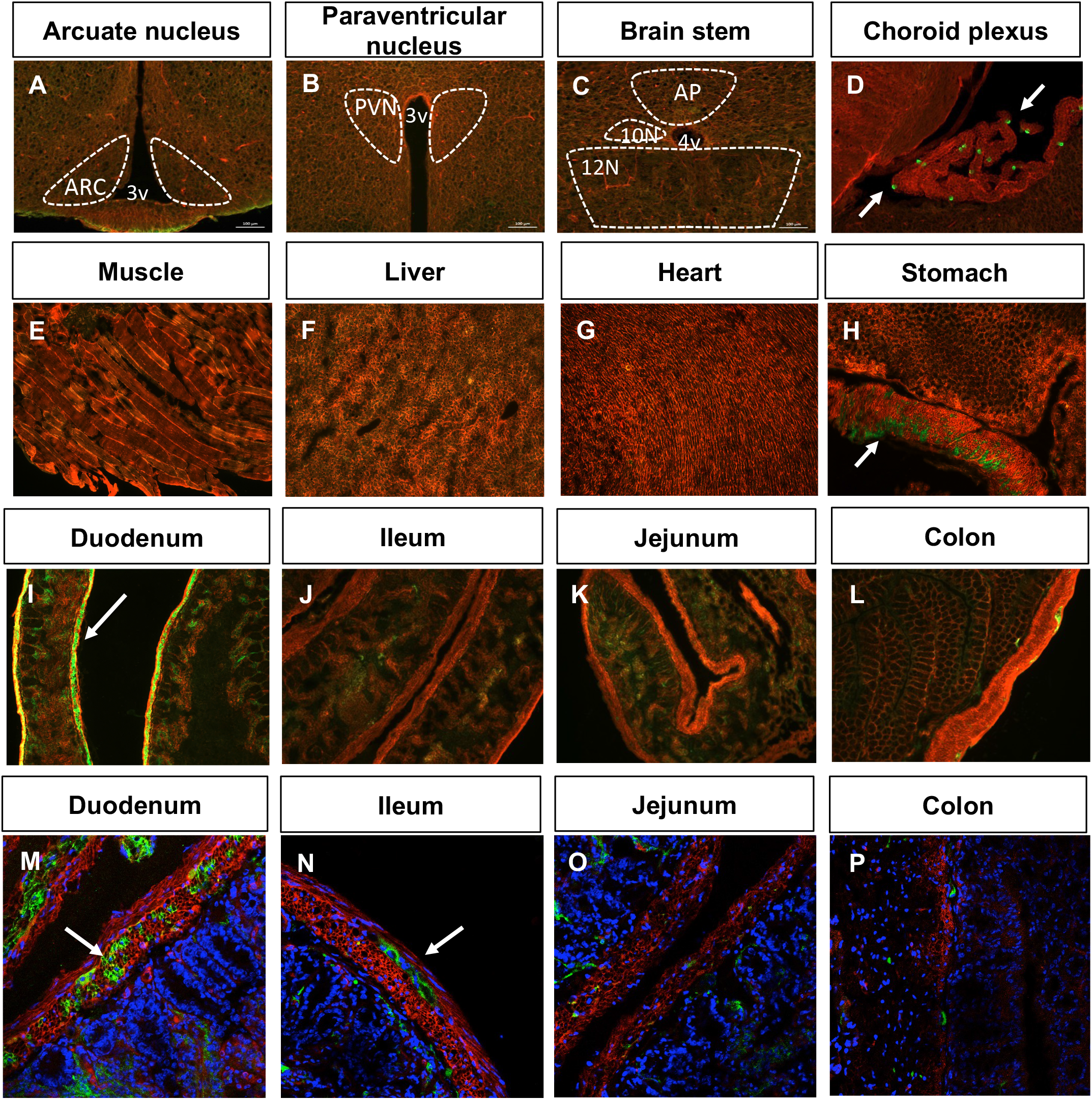
Cre-mediated recombination detected in the choroid plexus and duodenum of TetO-Cre^Jaw/J^ mice. Coronal sections of the brain of 18-20 week old TetO-Cre^Jaw/J^:mTmG male mice without doxycycline. *A*: Arcuate nucleus (ARC), *B*: paraventricular nucleus (PVN), *C*: brain stem, *D*: choroid plexus. 3v: third ventricle, 4v: fourth ventricle, AP: area postrema, 10N: dorsal motor nucleus of the vagus, 12N: hypoglossal nucleus. Sections of the *E*: muscle, *F*: liver, *G*: heart, *H*: stomach, *I*: duodenum, *J*: jejunum, *K*: ileum and the *L*: colon from 18-week old TetO-Cre^Jaw/J^:mTmG mice without doxycycline. Confocal images of *M*: duodenum, *N*: jejunum, *O*: ileum, *P*: colon from 18 week-old TetO-Cre^Jaw/J^:mTmG mice without doxycycline (representative images of *n* = 1-3 mice, male or female).

There was no evidence of spontaneous reporter transgene recombination in mTmG (TetO-Cre^Jaw/J^ negative) control mice (Supplemental Figure 1), demonstrating that green cells in TetO-Cre^Jaw/J^ positive mice were the result of Cre-recombinase expression in the absence of any tetracycline responsive transactivator. Low levels of GFP were detected in the intestine of 10-day old pups from TetO-Cre^Jaw/J^:mTmG mice, although this may also be auto-fluorescence of intestinal contents (Supplemental Figure 2), suggesting limited TetO-Cre leakiness in early life when receiving doxycycline via breast milk. Overall, our analysis revealed unregulated expression of Cre recombinase in the choroid plexus and digestive tract of adult TetO-Cre^Jaw/J^ positive mice.

### MIP-itTA mice are glucose intolerant due to insufficient insulin secretion

It was recently shown that common drugs used to control transgene expression can have off-target effects on mitochondrial and metabolic function^28^. Moreover, there is increasing evidence illustrating that overexpression of transgenes previously believed to be inert (e.g Cre-recombinase, GFP, transactivator) can have significant effects on cell function and viability^34,35,41-43^. Based on these concerns, we performed a series of *in vivo* experiments to investigate impact of the Tet-Off system components on glucose metabolism in males and females over time. Of note, we did not include mice with floxed genes in these experiments, thus phenotypes are all directly related to expression of TetO System components and not excision of any other gene.

Glucose tolerance, insulin sensitivity and glucose-stimulated insulin secretion were assessed at 13-15 weeks of age following a 5-week wash-out of doxycycline (Figure 3A). Male or female TetO-Cre^Jaw/J^ mice responded similar to wild-type (WT), age and sex-matched littermates in all parameters (Figures 3B-I). In contrast, we observed significant glucose intolerance in all mice of both sexes expressing the MIP-itTA transgene following an OGTT (Figure 3B-C) and AUC analysis (Figure 3D-E), regardless of Cre-recombinase expression. There were no significant differences in mouse weights or fasting blood glucose (18 hours or 4 hours) for any genotype within each sex (Supplemental Figures 3A-F), although there was a trend toward higher glycemia in female mice expressing the complete MIP-itTA:TetO-Cre system (Supplemental Figure 3D).

**Figure 3:**
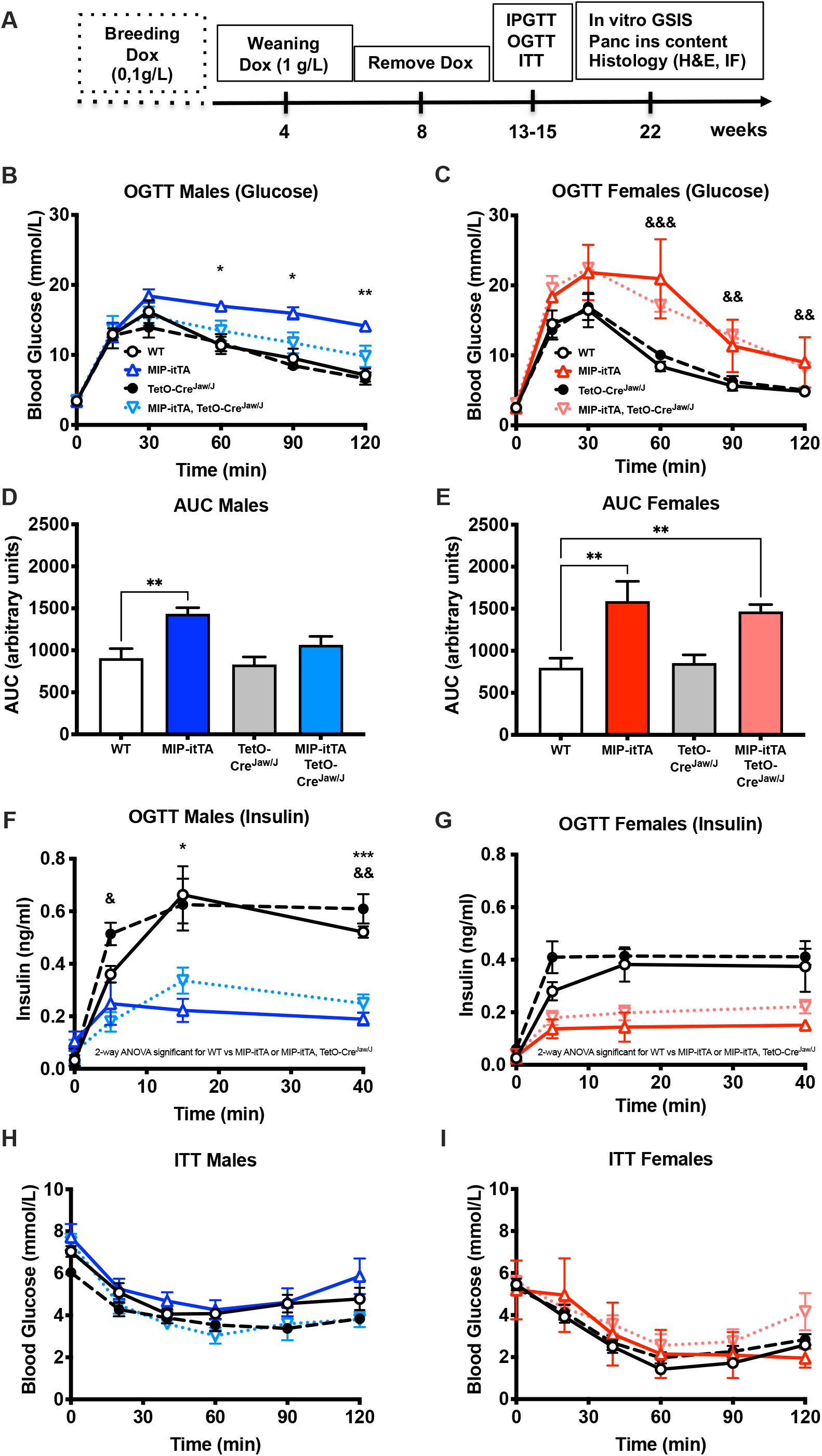
Mice carrying the MIP-itTA transgene are glucose intolerant and exhibit defective insulin secretion. *A*: Schematic of the experimental design and timeframe. Blood glucose following oral glucose challenge in 13-15 weeks old *B*: males (*n* = 4-6) and *C*: females (*n* = 2-5) and corresponding AUC (*D* for males and *E* for females) or serum insulin levels (*F* for males and *G* for females). *H-I*: Insulin tolerance test performed on mice from *B* and *C*. Post-hoc comparisons are shown for WT vs MIP-itTA (*p<0.05, **p<0.01, ***p<0.001) and WT vs MIP-itTA, TetO-Cre^Jaw/J^ (^&^p<0.05, ^&&^p<0.01, ^&&&^p<0.001). WT: white circle/solid line, MIP-itTA: open, red/dark blue triangle/solid line, TetO-Cre^Jaw/J^: black circle/dashed line, MIP-itTA, TetO-Cre^Jaw/J^: open, inverted pink/light blue triangle/dashed line.

To determine if glucose intolerance was due to defects in insulin secretion versus insulin action, we measured plasma insulin during the OGTT and noted that male and female mice carrying the MIP-itTA transgene also had a significant deficiency in glucose-stimulated insulin secretion (Figure 3F-G). In line with hyperglycemia of MIP-itTA+ mice being the result of a primary beta cell defect and not insulin resistance, all mice had similar responses to an injection of exogenous insulin (Figure 3H-I).

Finally, it is documented that mouse genetic background influences glucose metabolism^44-46^ and off-target effects of transactivator expression in the brain^35^. Doxycycline also has off-target effect depending on the mouse genetic background^47^. Initial groups of mice were a on mixed background of C57Bl/6N or 6J:129Sv:CD-1. To test whether the phenotypes were influenced by mouse strain or doxycycline, male mice were breed onto a pure C56Bl/6N background and glucose tolerance assessed by OGTT in mice that never received doxycycline. We again observed significant glucose intolerance correlating with presence of the MIP-itTA transgene (Supplemental Figure 3G-H) in the inbred mice. In short, we report that both male and female mice overexpressing the MIP-itTA are glucose intolerant due to a significant reduction in glucose-stimulated circulating insulin levels. This phenotype was independent of the TetO-Cre^Jaw/J^ transgene, which in our hands had no significant impact on glucose tolerance or GSIS *in vivo*.

### Tetracycline-controlled transactivators reduce insulin expression in beta cells

Reduced GSIS can result from low levels of insulin or defects in glucose sensing. Thus, we measured insulin content and secretion capacity of islets from these mice. As we saw identical GSIS defects in male and female mice, we performed subsequent experiments in male mice only. Islet architecture by H&E staining in pancreatic sections showed no visible differences between each genotype (Figure 4A-D). However, we noted insulin staining irregularities (e.g. high numbers of alpha cells in the islet core or cells that were neither insulin- or glucagon-positive) in MIP-itTA and MIP-itTA:TetO-Cre^Jaw/J^ islets (Figure 4E-H). Following analysis of more than 60 islets per genotype, we concluded that the presence of the itTA transgene correlated with having more than 30% of islets with these abnormal hormone staining patterns (Figure 4I). Consistent with these results, total pancreatic insulin content was reduced in MIP-itTA+ mice (Figure 4J) and primary islets from MIP-itTA or MIP-itTA:TetO-Cre^Jaw/J^ mice had significantly lower insulin content compared to littermate controls (Figure 4K). We also noted that the TetO-Cre^Jaw/J^ mice had slightly higher islet insulin content than the wildtype littermates (Figure 4K).

**Figure 4:**
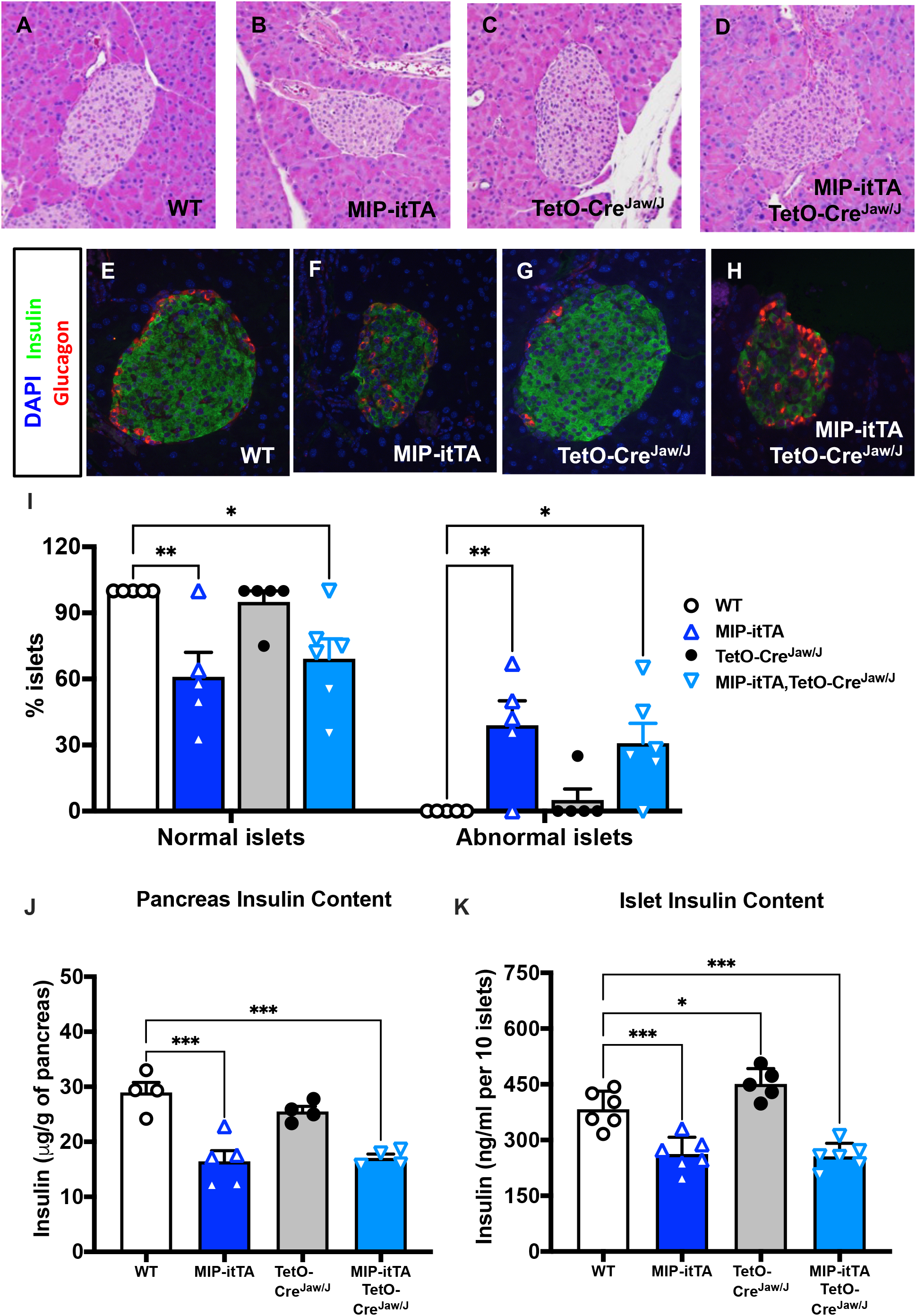
Islets from mice with the MIP-itTA transgene have abnormal insulin staining and decreased insulin content. Representative images of pancreatic islets of mice from each genotype following *A-D*: H&E or *E-H*: immunofluorescence staining with antibodies against insulin (green), glucagon (red), or DAPI (blue). *I*: Scoring of all the islets analyzed after immunofluorescence (*n* = 5-6 mice). *J*: Total pancreatic insulin content from males (*n* = 4-5). *K*: Insulin content of isolated islets from male mice from each genotype (pooled islets from *n* = 3 mice, values are means ± SD). Following ANOVA, post-hoc statistical comparisons are performed versus WT (*p<0.05, **p<0.01, ***p<0.001).

To determine if the defect in insulin content and secretion was the result of lower expression of insulin itself and/or other genes important for beta cell identity and function, we performed qPCR analysis on primary islets isolated from mice of each genotype. Correlating with reduced pancreatic and islet insulin content, mice harbouring the MIP-itTA transgene had reduced expression of both the Insulin 1 (*Ins1*) and Insulin 2 (*Ins2*) genes compared to the other control genotypes (Figure 5A). We also found that the gene for Solute Carrier Family 2 Member 2 (*Slc2a2* or *Glut2*), encoding a transporter that is the entry point for glucose into beta cells and plays a major role in glucose sensing, was significantly lower in the islets expressing the transactivator (Figure 5A). Genes for Pancreatic and Duodenal Homeobox 1 (*Pdx-1*), a transcription factor that controls pancreas development and beta cell identity, and Potassium Inwardly Rectifying Channel Subfamily J Member 11 (*Kcnj11*), encoding a voltage-gated potassium channel mediating hormone secretion, were not significantly different between genotypes (Figure 5A). Our results suggested that regardless of the presence of doxycycline, the tetracycline-controlled transactivator in itself may have detrimental effects on insulin expression.

**Figure 5:**
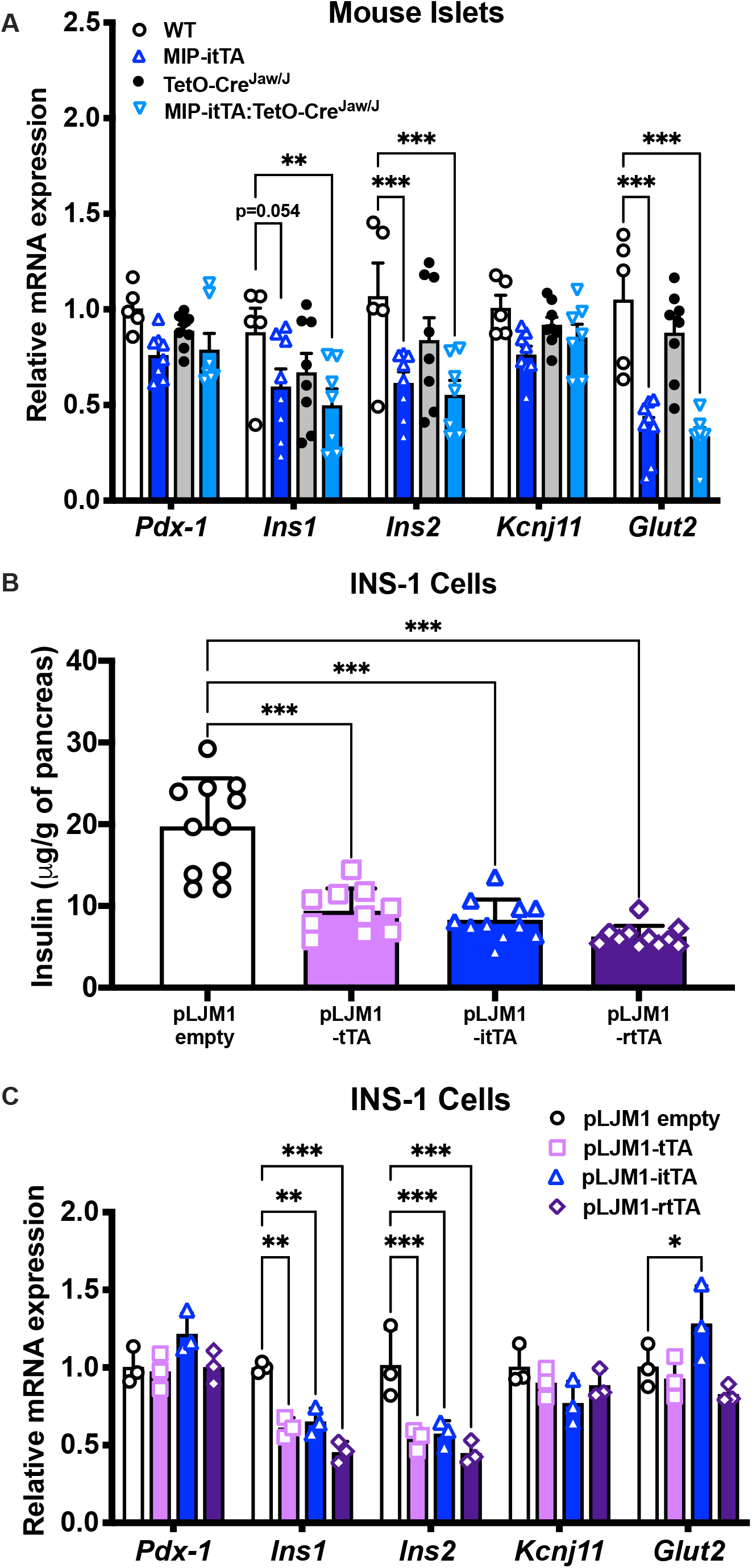
Expression of tetracycline-controlled transactivators in cultured beta cells reduces insulin mRNA and protein levels. *A*: Gene expression of primary islets of male mice expressed relative to *Tbp* (*n* = 5-8 mice per genotype, shown as means ± SEM. Comparison are versus WT following post-hoc analysis). *B*: Insulin content and *C*: gene-expression analysis of INS-1 cells stably overexpressing empty vector, tTA, itTA or rtTA. Values are means ± SD normalized to *Tbp* and representative of 3 independent experiments. *p<0.05, **p<0.01, ***p<0.001 represent comparisons to WT or pLJM1 empty, as indicated.

To address this hypothesis, we used lentiviral vectors to express cDNAs for either tTA, improved tTA (itTA) or reverse tTA (rtTA) under control of the CMV promoter in a rat insulinoma (INS-1) beta cell line. Following selection of stable clones, we measured insulin content and gene expression changes. Specific and similar expression of each transgene (tTA, itTA or rtTA) was confirmed across the clonal lines (Supplemental Figure 4). Consistent with data in mice and primary islets, insulin content in INS-1 cell lines expressing any of the tetracycline-controlled transactivators was decreased compared to levels in cells expressing empty vector (Figure 5B). We also observed a significant decrease in *Ins1* and *Ins2* gene expression following expression of the transactivators in this *in vitro* system (Figure 5C). Again, *Pdx-1* and *Kcnj11* were unchanged following transactivator expression (Figure 5C). In contrast, our stable cell lines did not have decreased *Slc2a2* (*Glut2*) gene expression and we noticed a significant increase with the itTA transgene. In summary, these data suggest that any tetracycline-controlled transactivator may cause a significant reduction in insulin expression leading to lower islet insulin content.

### Defects in insulin secretion linked to tetracycline-controlled transactivator expression were accentuated *in vivo*

Finally, as mice expressing the MIP-itTA construct had significantly deficient GSIS, we tested whether this functional defect was preserved *ex vivo*. Surprisingly, primary islets isolated from mice of each genotype had comparable insulin secretion in response to low (2.8 mM) or high (16.7 mM) glucose (Figure 6A), despite having lower insulin content (Figure 4K). However, response to KCl, a general stimulator of insulin secretion independent of glucose, was reduced in islets from all transgenic mice (Figure 6A). These data suggested that either primary islets contained sufficient stocks of insulin to respond effectively to the *in vitro* glucose challenge, or there were additional factors *in vivo* contributing to the defect in insulin secretion. As gut-derived hormones significantly enhance GSIS (known as the incretin effect^48^), an IPGTT was performed in males to test whether insulinopenia in MIP-itTA mice was due reduced incretin hormone action. We noted similar impairments in glucose intolerance following either i.p. injection (Figure 6B-C) as seen with oral gavage (Figure 3F-G) of glucose, suggesting that decreased GSIS in MIP-itTA+ mice was not the result of a defective incretin response. Taken together with the *in vitro* data, insulin reserve levels and additional factors *in vivo* likely exacerbate defects in GSIS linked to tetracycline-controlled transactivator expression in beta cells.

**Figure 6:**
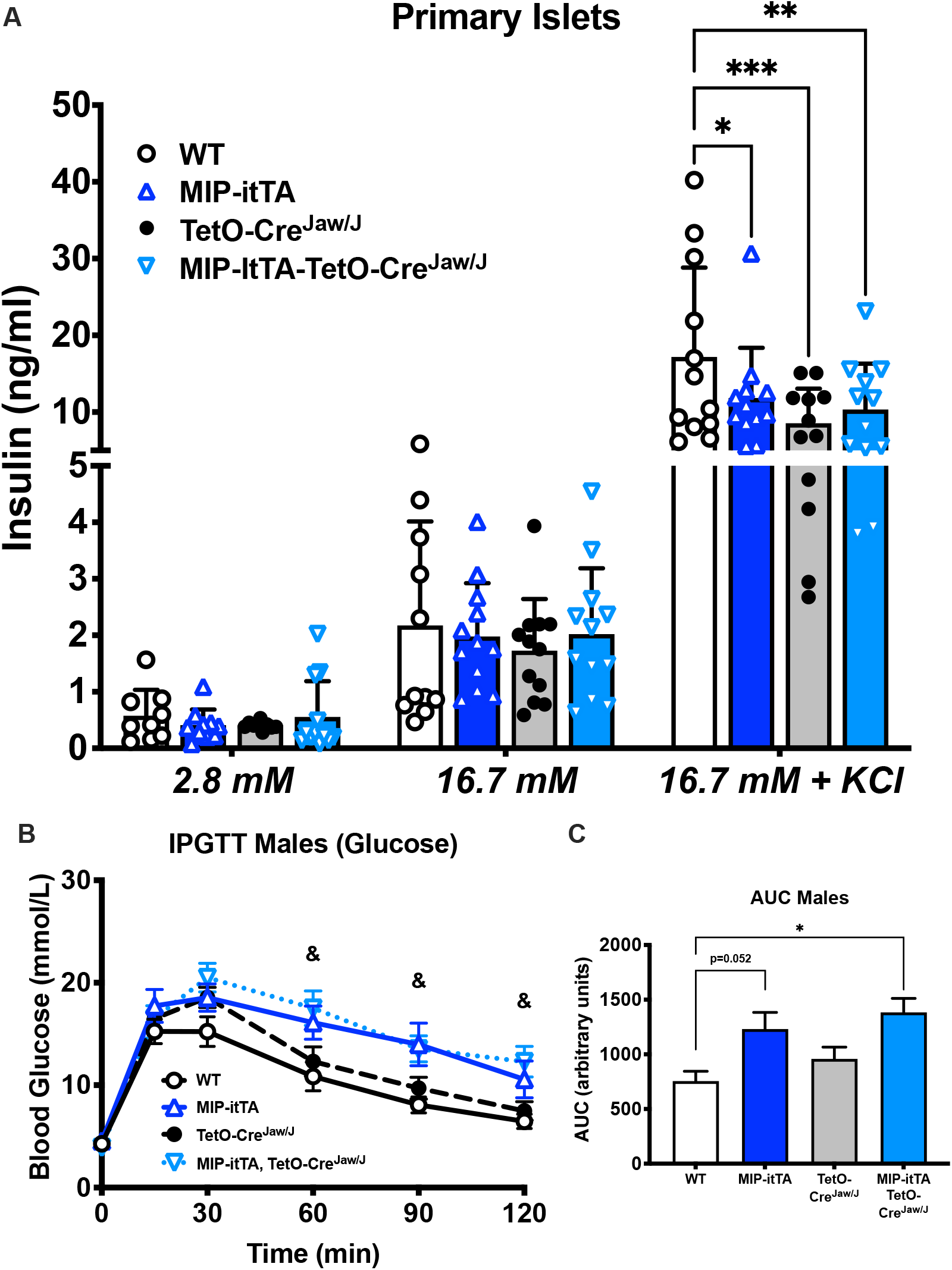
MIP-itTA expression decreases general beta cell secretory capacity. *A*: Glucose-stimulated insulin secretion from primary islets of male mice. Values are means ± SD of replicates of 10 islets from a pool of n=3 mice per genotype (*p<0.05, **p<0.01, ***p<0.001 versus WT). *B*: Blood glucose levels following intraperitoneal injection of glucose in 25 weeks old males (C57Bl/6N, never exposed to doxycycline). Values are means ± SEM. (*n* = 4-6 per genotype, (^&^p<0.05 comparing WT versus MIP-itTA, TetO-Cre^Jaw/J^). Area under the curve (AUC) of groups in *C*.

## DISCUSSION

Our work illustrates that a TetO-Cre/MIP-itTA approach effectively targets beta cells and allows efficient, drug-controllable expression of desired transgenes. This is in line with previously published reports using this approach^27,33,49^. However, our efforts reveal several caveats associated with these commonly used transgenic systems that could significantly affect interpretation of results produced in these models. We find that the TetO-Cre^Jaw/J^ transgene causes unregulated expression of Cre recombinase in the choroid plexus and regions of the gastrointestinal tract (stomach and duodenum). Moreover, we show that expression of a tetracycline-controlled transactivator in beta cells significantly reduces insulin expression, manifesting as defective glucose-stimulated insulin secretion, glucose intolerance and lower pancreatic insulin content in mice carrying a MIP-itTA transgene. Reduction in insulin content correlates with significant decreases in *Ins1/2* and *Glut2* mRNA expression in islets from transgenic mice that are reproduced following transactivator over-expression in cultured beta cells.

A large number of pancreas, islet and beta cell-specific transgene driver mouse lines are currently available to the scientific community. Over the last few decades, the beta cell field has struggled with tissue specificity of promoters and there are various examples of unexpected physiological and biological effects stemming from ectopic transgene expression^6,7,9,11^. These caveats are not limited to Cre-recombinase or beta cells, stimulating creation of a data repository to document off-target transgene activity in multiple lines of mice^50^. While extensively used and considered a highly controllable system, detailed data regarding leakiness or potential side effects of TetO and transactivator transgenes are not yet widely accessible. Availability of highly sensitive reporter mice (e.g. mTmG) greatly expands our ability to characterize these systems, facilitating identification of aberrant recombination events in even small and diverse subsets of cells. We identify cells of the choroid plexus, involved in the production of cerebrospinal fluid (CSF) and growth factors, and submucosal cells of the stomach and duodenal wall as areas where uncontrolled TetO-driven transgene expression could pose a problem. This could have undesired off-target effects depending on the transgene or gene target involved.

We chose to evaluate the Tet-Off approach in an effort to minimize toxicity and off-target effects caused by drug adminstration^28,29^. Tamoxifen can have long-term, negative impacts on function^17^ and proliferation of beta cells^23^, and doxycycline may alter mitochondrial function^29,51^. We wanted a model where drug administration was not required during the time of phenotypic or biochemical characterization. However, a drawback of the Tet-Off system is that doxycycline must be given to dams and pups during pregnancy and weaning, and then to adults up until the time of desired transgene expression. Long-term doxycycline exposure may have unexpected effects on progeny development^52,53^ and gut microbiota^54^ and experimental design should ensure that all mice, including controls, are exposed to similar experimental conditions including drug treatment.

The MIP-itTA/TetO-Cre system, in our hands, exhibited very low levels of leakiness in beta cells, which is expected and may not be a problem depending on the experiment. However, experimental designs requiring long-term maintenance of mice on doxycycline prior to transgene activation (i.e. induction desired during adulthood) could lead to accumulation of leaky recombination events in tissues that have low cell turnover, which includes beta cells. On the other hand, doxycycline can also accumulate in tissue depots, including the pancreas, maintaining effective levels of drug for several days/weeks after withdrawal^55^. This may necessitate longer wash-out periods to achieve 100% recombination efficiency. We observed Cre recombinase activity in about 70% of beta cells 4 weeks after doxycycline withdrawal; however, it is possible that a longer wash-out period would further increase efficiency.

In addition to limitations regarding recombination efficiency and drug exposure, caveats common to all systems, we noted significant impacts of the transactivator (tTA) proteins on glucose homeostasis. Effects of the MIP-itTA transgene alone on insulin levels and glucose tolerance were similar across sexes, but there was a small tendency toward a stronger effect in females (Figure 3C and Supplemental Figure 3D). Interestingly, our phenotype is very similar to results from mice with the tTA under the *Pdx1* promoter^56^. Nonetheless, our observations are in contrast to two other studies using tetracycline-controllable transactivators in combination with the *Ins1* promoter that report no effects of transactivator expression on glucose tolerance in male mice^27,49^. Differences in observations could be due to control groups used for comparisons, or other factors such as mouse strain or transgene-specific effects (e.g. level of transactivor expression). We found that altered glucose tolerance in the MIP-itTA mice was linked to defective GSIS and decreased pancreatic insulin content. Furthermore, we were able to reproduce reductions in beta cell insulin content and insulin expression *in vitro* by over-expression of either tetracycline-controlled transactivator (tTA), improved transactivator (itTA) or reverse transactivator (rtTA). Decreased *Insulin* mRNA expression was pronounced and reproducible in both primary islets and beta cell lines. Being transcriptional co-activators by design, it is possible that these foreign tTA proteins interact with and/or regulate transcription of these genes (and possibly many others) in unpredictable ways. We did not perform extensive analysis of other potential genomic targets or test whether the transactivators bind directly to the *Ins* gene promoters. However, our data suggest that off-target effects of tTA proteins are cell-autonomous, may be linked to dysregulation of the genome, and are shared by all of the tetracycline-controllable transactivator variants.

Despite significant effects of transactivator expression on insulin content and secretion *in vivo*, we were surprised to see no effect of transactivator expression on GSIS *in vitro*. However, this may be because beta cells produce insulin in excess, with only ∼5% released in response to a single stimulus^57^. Therefore, one glucose challenge may not be sufficient to reveal secretion deficiencies linked to low insulin content in primary islets *in vitro*, and repeated glucose stimulations may be necessary^58^. Depletion of insulin content might become more pertinent if these genetic tools are used in murine models of diabetes, where insulin resistance and beta cell stress lead to insulin hypersecretion and/or deficiency. Although we did not challenge our mice with obesogenic diets or beta cell stresses associated with diabetes, our data suggests that tetracycline-controlled transactivator expression has the potential to alter outcomes depending on the assay system.

Viable alternatives for inducible genome editing include tamoxifen-inducible systems or viral vectors expressing proteins under tissue-specific promoters, such as adeno-associated virus (AAV)8-Ins1-Cre^59^. These technologies also have limitations including technical difficulties to achieve 100% efficacy, beta cell toxicity and the potential for viral expression in other cell types. Surmounting evidence illustrates that there is no perfect system, but measures can be taken in all systems to control for expected and unexpected off-targets effects. Novel tools are continuously being developed. Self-cleaved inducible CreER (sCreER) may help to increase efficiency of recombination^60^ or a photoactivatable Cre^61^ could non-invasively and temporally activate transgene expression without the need for drugs. Of course, with new technology comes new caveats that will themselves require characterization and their own unique controls.

## Acknowledgements

Authors would like to thank Michel Fries for technical assistance with the lentivirus. Special thanks to Aurèle Besse-Patin, Émilie Courty, Cristina Bosoi and Preeti Bhatt for technical assistance and to Michel Cayouette, Katrina Podsypanina and Marie Kmita for reagents. JLE and NJ are the guarantor of this work and take responsibility for the contents of the article.

## Author Contributions

K.B. and T.A. processed the brains and analyzed corresponding immunofluorescence images. J.M. performed and analyzed whole-body pup staining. C.B. created expression constructs and lentiviral vectors and helped perform metabolic tests. N.J. contributed to the study design and conducted all other experiments. T.A. edited the manuscript. N.J. and J.L.E. designed the research, interpreted data, co-wrote and edited the manuscript. All authors gave final approval of the version to be published. N.J. and J.L.E. are the guarantors of this work and, as such, have full access to all data in the study and take responsibility for the integrity of the data and accuracy of data analysis.

## Prior presentation

This work has been presented at the virtual Diabetes Canada/CSEM Professional Conference 2020 as a poster (October 28-30, 2020).

## FIGURE LEGENDS

**Supplemental Table 1.**
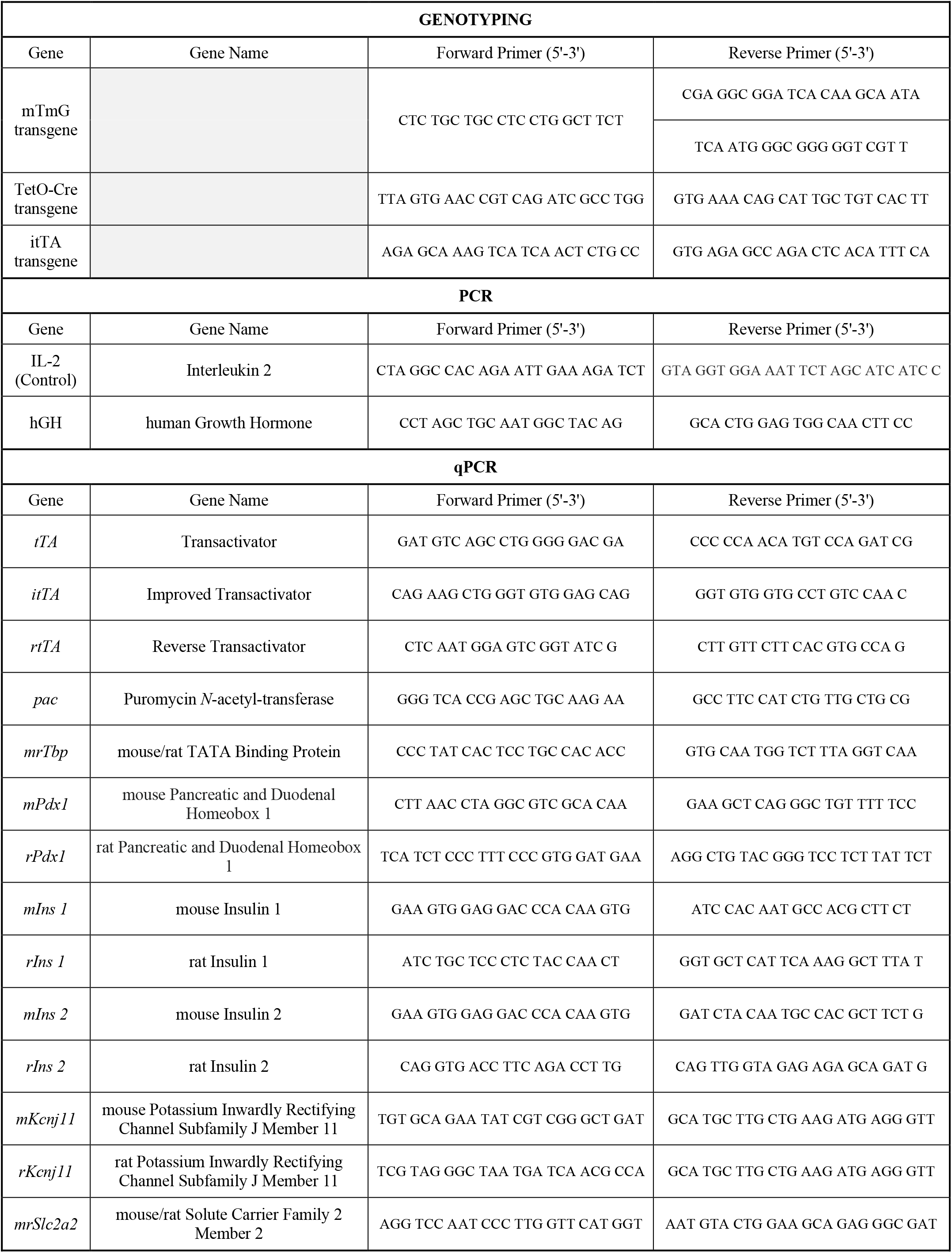

**Supplementary Figure 1:**
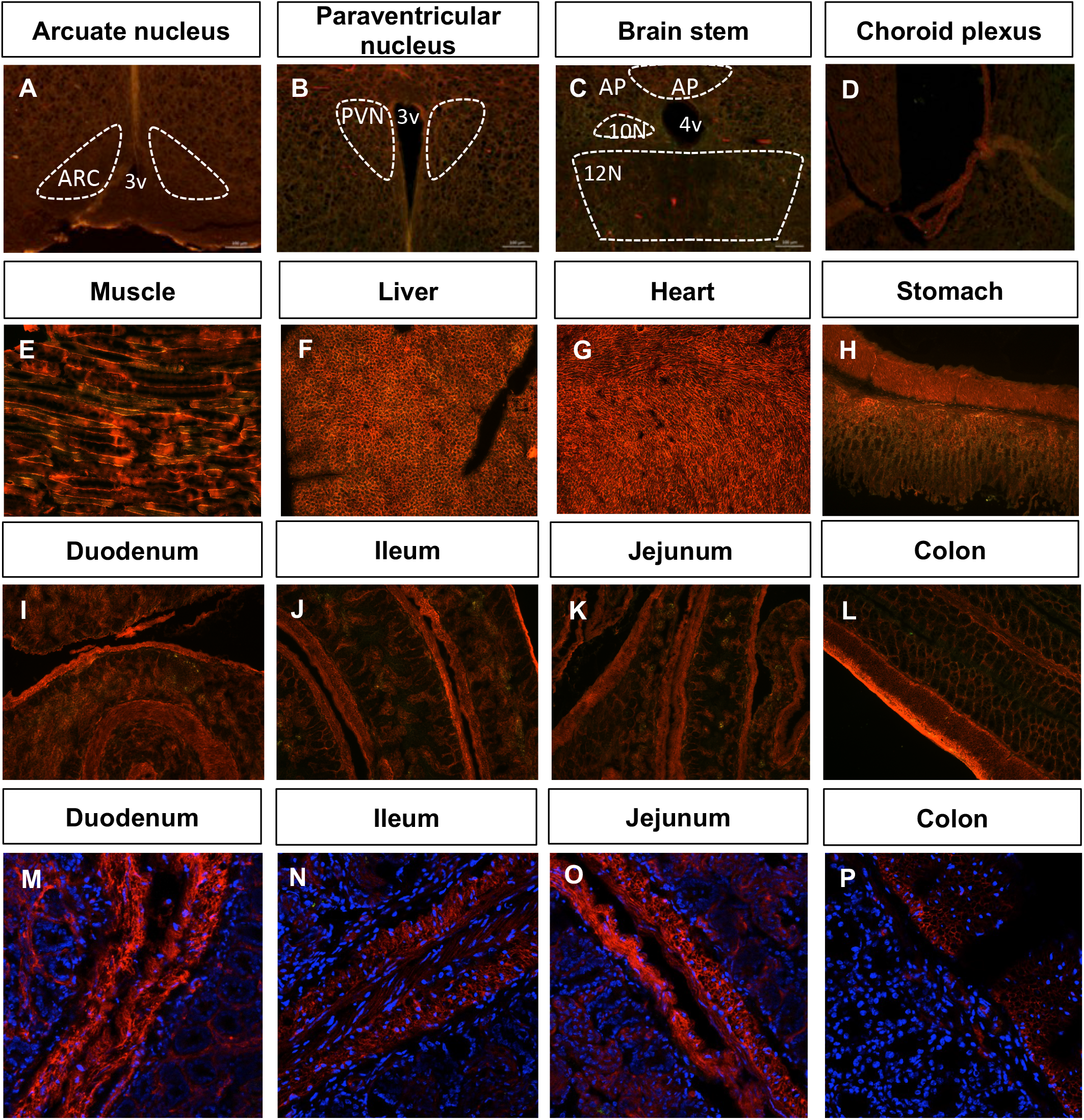
Cre-mediated recombination is not detected in mTmG (WT) littermate control mice. Coronal sections of brain from 18-20 week-old WT mTmG mice without doxycycline. **A)** Arcuate nucleus (ARC), **B)** paraventricular nucleus (PVN), **C)** brain stem, **D)** choroid plexus. 3v: third ventricle, 4v: fourth ventricle, AP: area postrema, 10N: dorsal motor nucleus of the vagus, 12N: hypoglossal nucleus. Sections of the **E)** stomach, **F**) muscle, **G**) liver, **H**) heart, **I)** duodenum, **J)** jejunum, **K)** ileum and the **L)** colon from 18 week-old WT mTmG mice without doxycycline. Higher magnification, confocal images of **M**) duodenum, **N**) jejunum, **O**) ileum, **P**) colon from 18 week-old TetO-Cre^Jaw/J^:mTmG mice without doxycycline (representative images of *n* = 1-3 male or female mice).

**Supplementary Figure 2:**
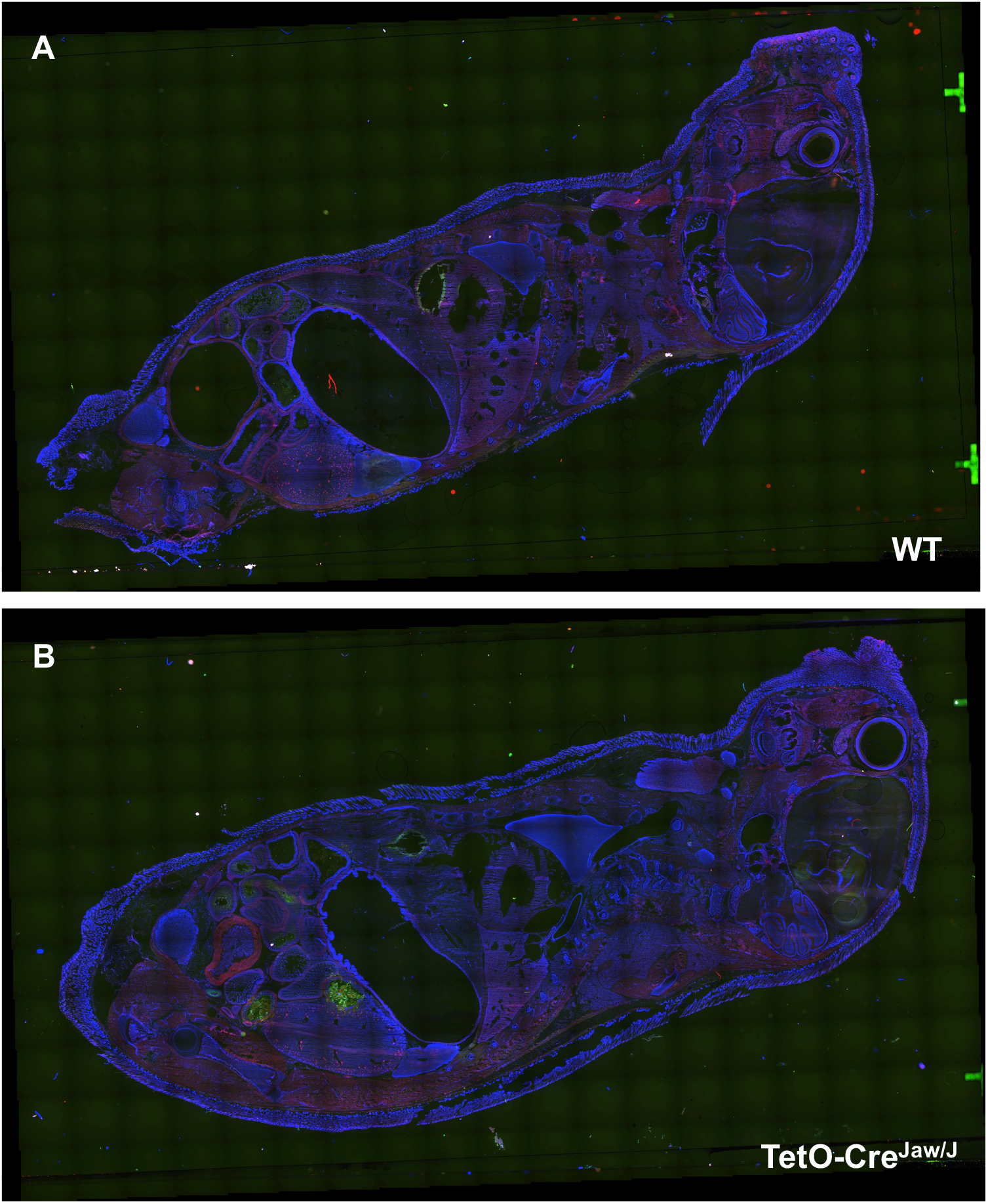
Cre-mediated recombination is not detected in 10-day old pups from WT or TetO-Cre+ mice. Immunofluoresecence images of whole-mount 10 day old pups from **A)** mTmG or **B)** TetO-Cre^Jaw/J^:mTmG mice that were never exposed to doxycycline.

**Supplementary Figure 3:**
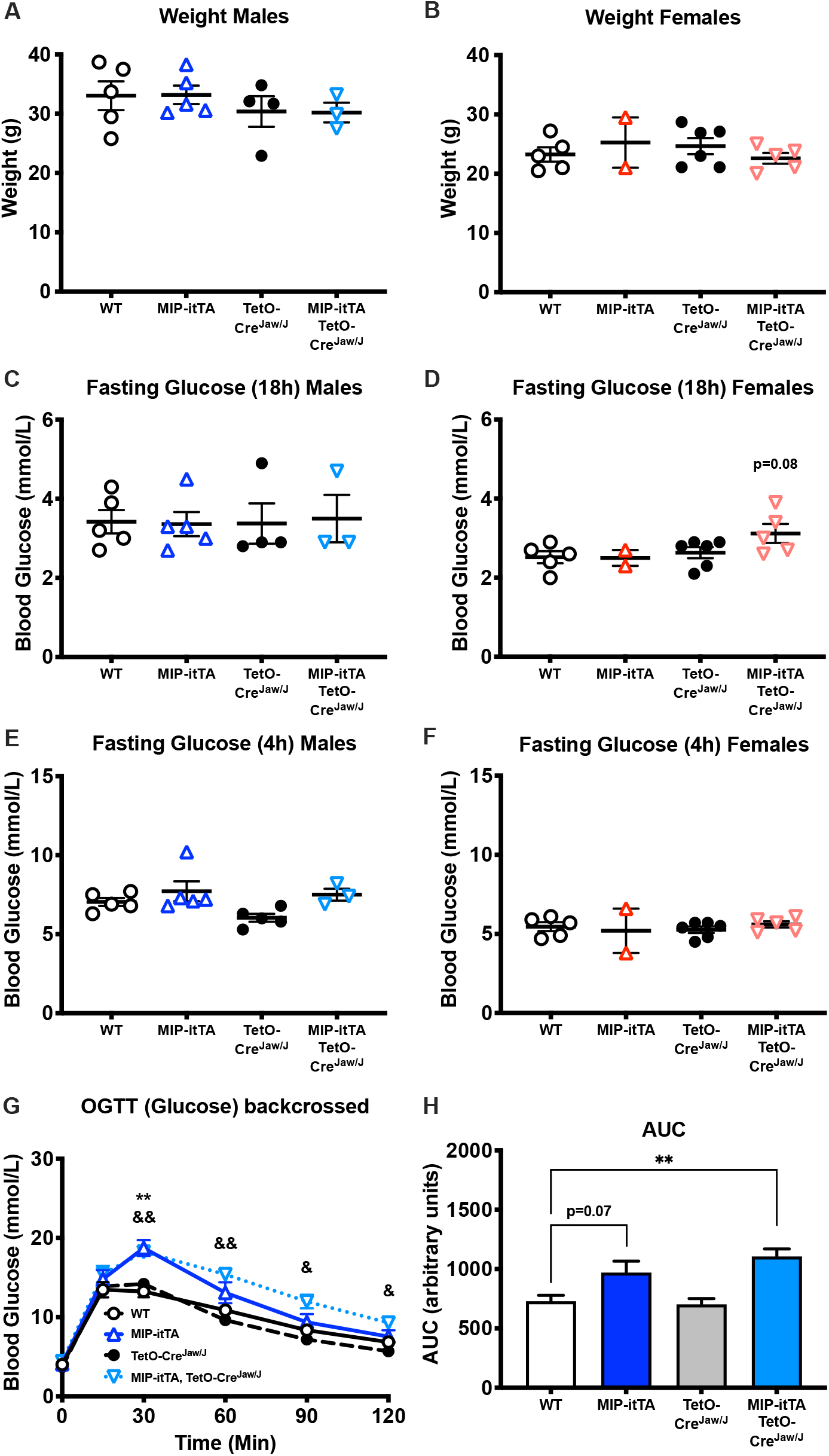
Transgene expression has no effect on weight or fasting blood glucose, while glucose tolerance is decreased by MIP-itTA. Body weights of 13-15 week old **A)** males (*n* = 4-6) and **B**) females (*n* = 2-5). **C**,**D**) 18 hour or **E**,**F**) 8 hour fasting blood glucose levels in males or female mice, as indicated. **G**) Blood glucose following oral glucose challenge in 25 week old males, never exposed to doxycycline, on a pure C57Bl/6N background (*n* = 4-6). **H**) Area under the curve for groups in **G**). WT: white circle/solid line, MIP-itTA: open, dark blue triangle/solid line, TetO-Cre^Jaw/J^: black circle/dashed line, MIP-itTA, TetO-Cre^Jaw/J^: open, inverted light blue triangle/dash line. Post-hoc comparisons are shown for WT vs MIP-itTA (**p<0.01), and WT vs MIP-itTA:TetO-Cre^Jaw/J^ (^&^p<0.05, ^&&^p<0.01).

**Supplementary Figure 4:**
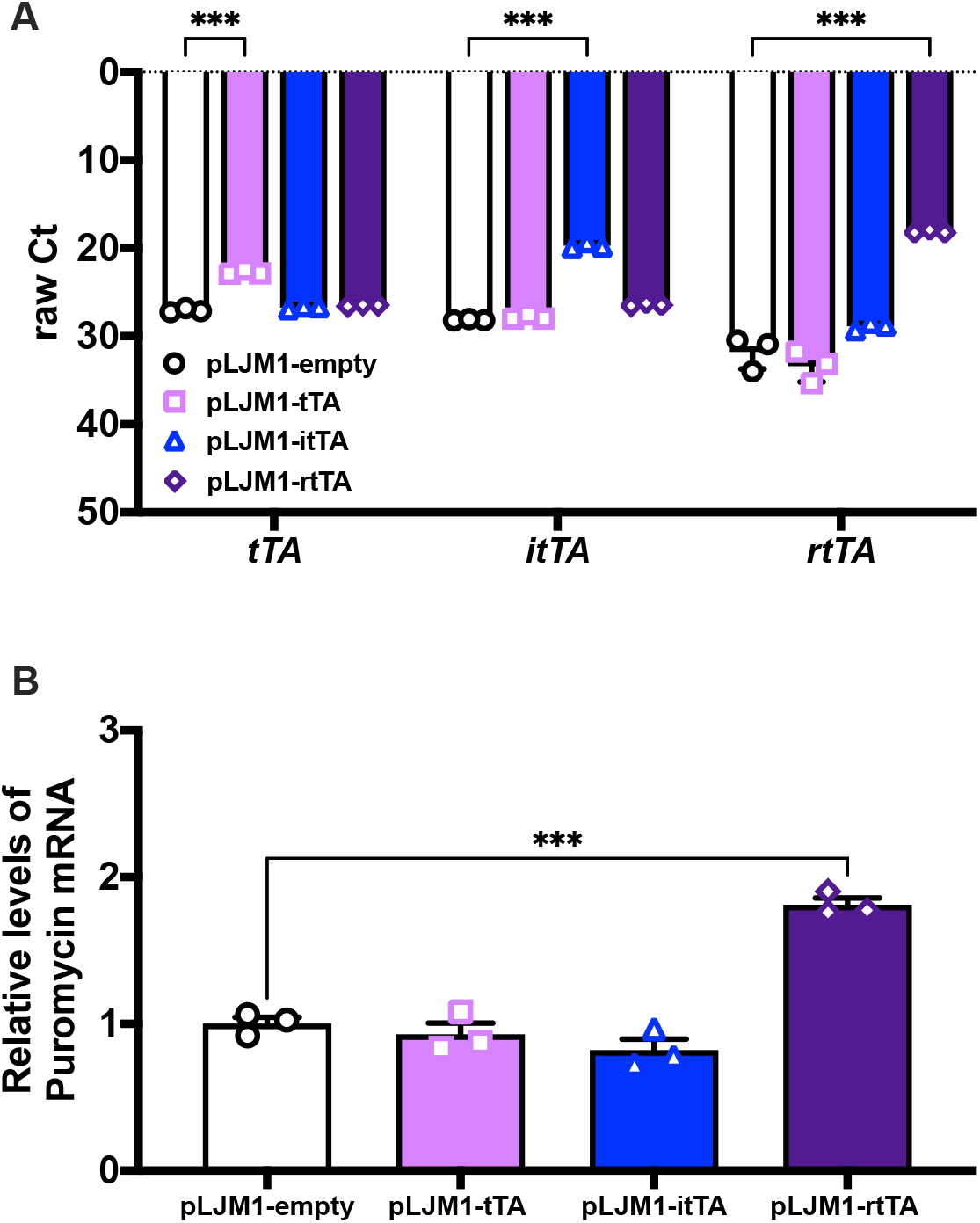
Relative expression levels of the tetracycline-controlled transactivators stably expressed in INS-1 cells. **A)** Raw Ct values for each transactivator measured by pPCR from mRNA isolated from each stable cell line. Given poor sequences similarity between cDNAs, individual primer sets were required for each construct. **B)** Relative mRNA expression of the puromycin gene, common to the expression vector used to make the stable cell lines. Values are means ± SD of biological triplicates, with statistical comparisons made to pLJM1 empty vector (***p<0.001).

